# Coordination among leaf and fine root traits across a strong natural soil fertility gradient

**DOI:** 10.1101/2023.11.02.565353

**Authors:** Xavier Guilbeault-Mayers, Hans Lambers, Etienne Laliberté

## Abstract

Unravelling how fundamental axes of trait variation correlate among leaves and roots and relate to nutrient availability is crucial for understanding plant distribution. While the leaf trait variation axis is linked to nutrient availability gradients, the response of root trait variation to the same gradients yields inconsistent results.

We studied leaf and root trait variation among 23 co-occurring plant species along a 2-million year soil chronosequence to assess how leaf and root traits coordinate and how this resulting joint axis of variation relates to soil fertility.

Mycorrhizal association types primarily structured the axes of leaf and root trait variation. However, when considering species abundance, soil nutrient availability was an important driver of trait distribution. Leaves that support rapid growth in younger, fertile soils were associated with roots of larger diameter and arbuscular mycorrhizal colonization. In contrast, leaves that favour nutrient conservation in nutrient-impoverished soil were associated with greater root hair length and phosphorus-mobilizing root exudates.

At the species level, the signals deviated from the community-wide results presented above, highlighting the challenge of generalizing a specific set of root trait values that consistently meet the requirements of leaves supporting either rapid growth or survival.

## Introduction

Evaluating how fundamental axes of trait variation - describing strategies ranging from those supporting rapid growth to survival - coordinate between leaves, influencing nutrient use, and fine roots, influencing nutrient acquisition, is crucial for enhancing our understanding of plant functioning and distribution along environmental gradients, including soil fertility (Reich, 2014). A growing number of plant species in global trait databases (Wright et al., 2004; Bergmann et al., 2020) and root trait measurements such as root exudates (reviewed in Wen et al., 2022) have improved our understanding of interspecific variation in leaf and root traits across biomes. However, the general lack of environmental metadata (e.g., soil fertility) in these databases leaves uncertainties about leaf and root trait covariation and their coordination with soil variables (Weigelt et al., 2021). Because soil nutrient availability might impact fine root traits that influence nutrient acquisition differently from leaf traits that influence nutrient use (Weemstra et al., 2016), more studies of root and leaf trait covariation across soil nutrient availability gradients are needed.

According to nutrient economic theory, plant performance depends on leaf trait values that support rapid growth where nutrients are abundantly available (e.g., high specific leaf area, SLA), or slow growth and conservation of acquired nutrients such as nitrogen (N) and phosphorus (P) allowing higher long-term nutrient-use efficiency under adverse conditions (e.g., high leaf tissue density, LTD; Craine, 2009). This trade-off has been reported widely in different studies on leaf traits, across and within biomes (e.g., Reich et al., 1992; Wright et al., 2004). Leaves that promote rapid growth enhance carbon (C) assimilation by minimizing both C investment and C maintenance costs associated with extended longevity but incur greater nutrient losses during leaf senescence than conservative leaves do, owing to their high nutrient investment in short-lived leaves that exhibit less efficient nutrient remobilization. Conversely, long-lived conservative leaves enhance nutrient-use efficiency by allocating more C relative to N and P per leaf area and mass, resulting in a well-defended leaf that reduces nutrient loss, but limits the C assimilation rate (Wright et al., 2004). This range of strategies is termed the leaf economics spectrum (LES) or referred to, in terms of return on C investment, as the ‘fast-slow’ spectrum (Wright et al., 2004).

A spectrum similar to the LES was anticipated within the root economic space (RES) to enable rapid growth or promote nutrient conservation (Reich, 2014), which was later identified as the conservation gradient (i.e., root nitrogen concentration [N]-root tissue density (RTD) axis) (Bergmann et al., 2020). Analogous to the LES, this spectrum impacts root nutrient-acquisition rate, lifespan and nutrient loss relative to C investment (Bergmann et al. 2020). However, since soil nutrients are less mobile than aboveground resources (e.g., CO_2_), plants must forage soil nutrients (Laliberté, 2017). Consequently, the RES is multidimensional, including another root trait axis, the collaboration gradient, describing a range of nutrient-foraging strategies (Bergmann et al. 2020), enabling plants to alleviate limitations associated with low nutrient mobility. Along this gradient (i.e., diameter-specific root length (SRL) axis), for a given root mass, fine roots may forage for nutrient by increase their length producing a root of high SRL or rely on an increased diameter to favor arbuscular mycorrhizal colonization to forage nutrient beyond the root influence zone (Bergmann et al., 2020). However, the variation in root traits does not appear to be clearly connected to fast growth or survival, as they are not consistently related to variation in soil nutrient availability. This creates uncertainty about how the RES coordinates with the LES (Weemstra et al., 2016), and whether a single ‘fast-slow’ spectrum (Reich, 2014) is a valid concept for describing the joint variation of root and leaf traits.

Theoretically, rapid growth should be associated with leaf and root traits maximizing C gains, suggesting a coordination among both ‘fast’ ends of the LES and conservation gradient and fertile soil. This framework generates a single ‘fast-slow’ spectrum where variation along the diameter-SRL axis should be unrelated to nutrient availability gradient. Meanwhile, trait values along the root [N]-RTD axis should vary with this gradient, thereby forming a ‘fast’ and ‘slow’ RES subspace consistently favored in fertile and infertile soils, respectively. This aligns with the fact that the majority of studies investigating leaf and root trait coordination have reported a positive root [N] - leaf [N] correlation (Weigelt et al., 2021). It is also in line with the observation that ‘fast’ leaf traits are consistently associated with more fertile soils at a global scale (Ordoñez et al., 2009), while the diameter-SRL axis exhibits contrasting responses to nutrient availability (Table S1). Among 17 studies, 10 showed increasing root diameter and decreasing SRL under higher fertility and seven found the opposite. However, high root [N] has also been associated with nutrient-impoverished soil, enabling rapid nutrient acquisition to compete with microorganisms (Liu et al., 2010; Freschet et al., 2017), and required in species adapted to these soils to rapidly exude carboxylates, providing access to recalcitrant P (Wen et al., 2022). These findings challenge the validity of a single ‘fast-slow’ spectrum, as the ‘fast’ subspace of the RES may be linked to ‘slow’ leaf trait syndrome in nutrient-poor soil. Furthermore, traits directly related to the use of organic and recalcitrant nutrients and factors that influence root trait variation are often overlooked. These include the extent of investment in ectomycorrhizal and ericoid symbiosis (Smith & Read, 2008), carboxylate exudation (Wen et al., 2022), growth forms (Ma et al., 2018), and mycorrhizal association types (Valverde-Barrantes et al., 2017). Overall, this introduces uncertainty regarding the primary drivers of root trait variation and if a single ‘fast-slow’ spectrum is a valid concept for predicting covariation between leaf and root traits, where nutrient availability may represent the primary source of environmental variation.

To evaluate how fine root and leaf trait covariation might depend on soil fertility, we conducted our study along a long-term soil chronosequence in Western Australia (Turner et al., 2018). Long-term soil chronosequences are series of adjacent soils that differ in age but are formed under similar environmental conditions from the same parent material (Walker et al., 2010). Large differences in soil age give rise to a strong natural fertility gradient and minimize plant responses to other environmental factors (e.g., climate, topography, parent material) (Walker et al., 2010). Along the Warren chronosequence (Turner et al., 2018), we measured common leaf and root traits of 23 of the most abundant plant species including one very common species, *Agonis flexuosa*, which is one of the most abundant species in all stages of the chronosequence. We also measured foliar manganese concentrations [Mn] that can be used to proxy rhizosphere carboxylate concentrations, since Mn is mobilized by carboxylates and stored in leaf vacuoles (Lambers et al., 2015a). Our objectives were to 1) determine whether soil nutrient availability has a stronger influence on root trait variation than growth form or mycorrhizal symbiosis type, and 2) determine how leaf and fine-root traits covary along a soil fertility gradient.

## Materials and methods

### Study site

Sampling took place from September to December 2016 along the Warren dune chronosequence, located in d’Entrecasteaux National Park in Western Australia. The plant communities studied are located at five stages whose ages are known approximately and whose soils range from unstable sand, leached calcareous sand, podzol over calcareous sand and podzol on siliceous sand (Turner et al., 2018). Turner et al. (2018) posit that the ages of the dunes range from the Early Pleistocene to the Holocene as dune systems on the Western Australian coastal plain are linked to sea level changes during interglacial periods throughout the Pleistocene (Playford et al., 1976; Kendrick et al., 1991). Each major dune system in the chronosequence corresponds to a stage of fertility (Turner et al., 2018). For reference, the dune systems under study correspond to dune stages 1, 2, 3, 4 and 6 in the study by Turner et al. (2018). Within each of these stages, five 20 m x 20 m plots were randomly positioned. As shown by Turner et al. (2018), variation in soil P and N availability corresponds to the soil formation model developed by Walker & Syers (1976). Topsoil (0-20 cm) concentrations of nutrients such as total P, total N and ‘readily-available’ P where roots were sampled, however, did not show the same patterns as the first 1 m of soil. Plant nutrient uplift likely increased soil nutrient concentration in stages 2 through 4, peaking at stage 3 (Table S2) (Turner et al., 2018). There is no significant difference in temperature and precipitation across the chronosequence over the relatively short distance (∼10 km) from the youngest to the oldest dunes (Turner et al., 2018). The climate is Mediterranean, with hot, dry summers and mild, rainy winters, with a mean annual temperature and precipitation of 15.2°C and 1185 mm, respectively (Turner et al. 2018).

### Soil chemical analyses

Four soil samples, one in each quarter of the plot, were collected from the first 20 cm of the mineral soil in each plot. Samples were then combined into a composite sample, which was sieved (< 2 mm) and air-dried before conducting soil chemical analyses at the Smithsonian Tropical Research Institute (Turner et al., 2018). Nitrogen was determined by dry combustion using a Thermo Flash 1112 analyzer (Thermo Fisher Scientific, Massachusetts, United States). Total P concentration was determined by ignition at 550°C for 1 h followed by acid extraction with 1 M H_2_SO_4_ for 16 h. Determination of exchangeable P (resin P) was carried out using anion exchange membranes (Turner & Romero, 2009).

### Leaf and root sampling

Roots and leaves from multiple individuals of selected plant species were sampled within each plot to obtain a composite sample per species. In total, 111 aggregate samples were collected, and among these samples, strategies and growth form had a relatively constant representation (Tables S3-S6). Leaves directly exposed to light and fine roots were sampled from the most abundant species that together represented a relative canopy coverage as close as possible of 75% (Table S3). Roots were collected in the first 20 cm of the soil layer directly from targeted individuals. The nutrient-acquisition strategies for all composite samples were assigned based on a threshold of 10% of symbiotic root colonization, according to the overall apparent rate of misdiagnoses for mycorrhizal structures (Brundrett, 2009), except for ECM and ERM colonization, for which we were confident of the diagnosis.

### Leaf functional traits

Leaves were photographed in the field using a Portable Imaging & Calibration Kit (PICK, Régent Instruments Inc, Quebec, Canada), Olympus TG-4 tripod cameras (Olympus corporation, Tokyo, Japan) and subsequently analyzed using WinFolia software (Régent Instruments Inc, Quebec, Canada) to determine leaf area. Leaf thickness was also measured in the field using a digital caliper (Sona Enterprise, California, United States). Following this, leaves were dried on site and re-dried in the laboratory at 60°C for 72 h and weighed to obtain the specific leaf area and leaf tissue density. Leaf [N] was obtained using an elemental analyzer (Vario Micro Elementar, Langenselbold, Germany). Leaf [P] and [Mn] were determined after acid digestion of 2 mg of dry leaves followed by an inductively coupled plasma mass spectrometry analysis (Perkin Elmer NexION 300x, Waltham, USA).

### Root functional traits

Root functional traits were measured on fine roots that were classified using a root order-based threshold. For each species, variation in color, texture, diameter and rigidity were used to distinguish fine roots, resulting typically in the first two to three root orders being sampled. Specific root length, root diameter, root branching intensity (BrInt) and root tissue density were obtained using WinRhizo Pro software (Régent Instruments Inc, Quebec, Canada). Dry mass was obtained by drying fine roots at 60°C for 72 h to complete the measurement of SRL and RTD. Root hair density (RHD) was measured on 10 segments of approximately 1 mm across 10 randomly sampled first-order roots and root hair length (RHL) was measured on one randomly sampled root hair per first-order root previously sampled for RHD using images captured with a Zeiss Axio Imager 2 microscope (software: AxioVison, Jena, Germany).

### Degree of mycorrhizal colonization

Fine roots were cut into sections of approximately 1 cm length and then bleached in modified syringes (Claassen & Zasoski, 1992) using 10% (v/v) KOH at 90°C. Cleared roots were acidified with 5% (v/v) acetic acid for five minutes. Subsequently, roots were stained in Shaeffer Black ink and 5% (v/v) vinegar solution for 4 min (Vierheilig et al., 1998). Afterwards, roots were kept in a lactoglycerol solution for 48 h to remove excess staining. Finally, roots were mounted on a microscope slide with glycerol and observed at X100 for an overview and at a higher magnification up to X400 for identification of fungal structures.

To derive a continuous variable of the degree of arbuscular and ericoid mycorrhizal colonization, approximately 900 mm of fine roots were selected, and 300 mm of these root fragments were systematically sampled and analyzed per segments of approximately 0.15 mm using a similar methodology as proposed by Trouvelot (1986). The degree of colonization of each segment which considered all fungal structures (i.e., arbuscules and hyphae), was determined based on seven classes (1: 0%; 2: <1%; 3: <10%; 4: <25%; 5: <50%; 6: >50%; 7: >75%). The degree of colonization was obtained by summing the product of the median value of class *i* by the length of the segments associated with this class standardized by the total length of the analyzed segments:

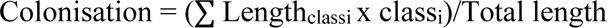

Ectomycorrhizal colonization was obtained by dividing the number of colonized root tips by the total number of root tips (20 tips on average per sample).

### Statistical analyses

All statistical analyses were performed in the R environment (R Core Team, 2020). Principal component analysis (PCA) was performed on standardized trait values, using the rda() function from the ‘vegan’ package (Oksanen et al., 2020). The significance of the PCA axes was assessed by permuting the data and comparing the observed inertia values of the axes with those obtained by chance, while trait loading significances were assessed using the broken-stick criterion (Peres-Neto et al., 2003). A PERMANOVA analysis was conducted on standardized traits and Euclidean distance, utilizing chronosequence stage, nutrient-acquisition strategy, and growth form as factors to evaluate which factors explained the greatest variance in RES and LES using the adonis() function from the ‘vegan’ package (Oksanen et al., 2020). To prevent circularity, symbiotic traits were excluded from the analysis, as they were initially used to define the nutrient-acquisition strategy. To evaluate the association between functional traits, species abundance and soil fertility, fourth-corner analysis was performed using the fourthcorner() function from ‘ade4’ package (Dray & Dufour, 2007). Model 2 permutation was used within the fourth-corner analysis where only rows of the abundance matrix were permuted, and false discovery rate was used for multiple comparison correction. Co-inertia analysis was used to assess the degree of covariation and correlation between the LES and RES using dudi.pca() and coinertia() function from ‘ade4’ package (Dray & Dufour, 2007) on unweighted and weighted traits by relative cover (ΣPiTi). The shared inertia (RV) coefficient between two multivariate tables, derived from co-inertia analysis, is high when both structures vary along a similar axis regardless of direction, and low when they vary independently (Dray et al., 2003). To test if the observed RV coefficient significantly differed from the RV coefficient distribution under the null hypothesis, permutations using the randtest() function from the ’ade4’ R package (Dray & Dufour, 2007) were used. To assess whether community-weighted mean traits reflected trait selection, not just variation in community composition, we compared trait values of common species with those of community-weighted means. Differences in strategies among stage classes were assessed with linear models (lm), generalized linear models (glm) from the ‘stats’ package (R Core Team, 2020) and generalized least squares (gls) from the ‘nlme’ package (Pinheiro et al., 2020). Models were modified with an appropriate variance function or appropriate family distribution to minimize heteroscedasticity and maximize normality of models’ residuals and were selected based on visual inspection of residual distributions and Akaike information criterion (AIC) (Zuur et al., 2009; Zuur et al., 2010). Post-hoc Tukey HSD tests were conducted using the ‘emmeans’ (Lenth, 2020) and ‘multcomp’ (Hothorn et al., 2008) packages. No linear analysis was performed on community-weighted mean values, since they were performed in the fourth-corner analysis. Finally, we also presented species leaf and root trait syndromes from the stages with the most contrasting nutrient availability (stage 3 and 5) to illustrate the range of strategies within community-wide trait syndromes.

## Results

### Root and leaf trait spectrum

Principal component analysis (PCA) performed on root traits revealed three independent axes of root trait variation. The first PCA axis, which we refer to as the root collaboration gradient, explaining 32.0% of the variance, ranged from species outsourcing nutrient acquisition through a large root diameter, high AM colonization and to a lesser extent ECM colonization to species with high SRL, BrInt, RHL and RHD values but low AM and ECM colonization intensities (Fig. 1a). The second axis of the PCA, explaining 20.4% of the variance, was more difficult to identify because root [N] was not measured and RTD contributed greatly to the first, second and third axes (Figs. 1a and S1a; Table S7). This pattern of correlation was mainly driven by ERM and ECM species, as they exhibited roots of high tissue density (Fig. S2b). Our analysis also highlighted a third significant dimension representing 15.1% of root trait variation (Figs 1a and S1a). The contribution to the third dimension was also mainly driven by ECM and ERM colonization and their coordinating traits, since only these symbiotic traits as well as branching intensity and root tissue density were involved (Fig. 1a; Table S7).

**Figure 1:**
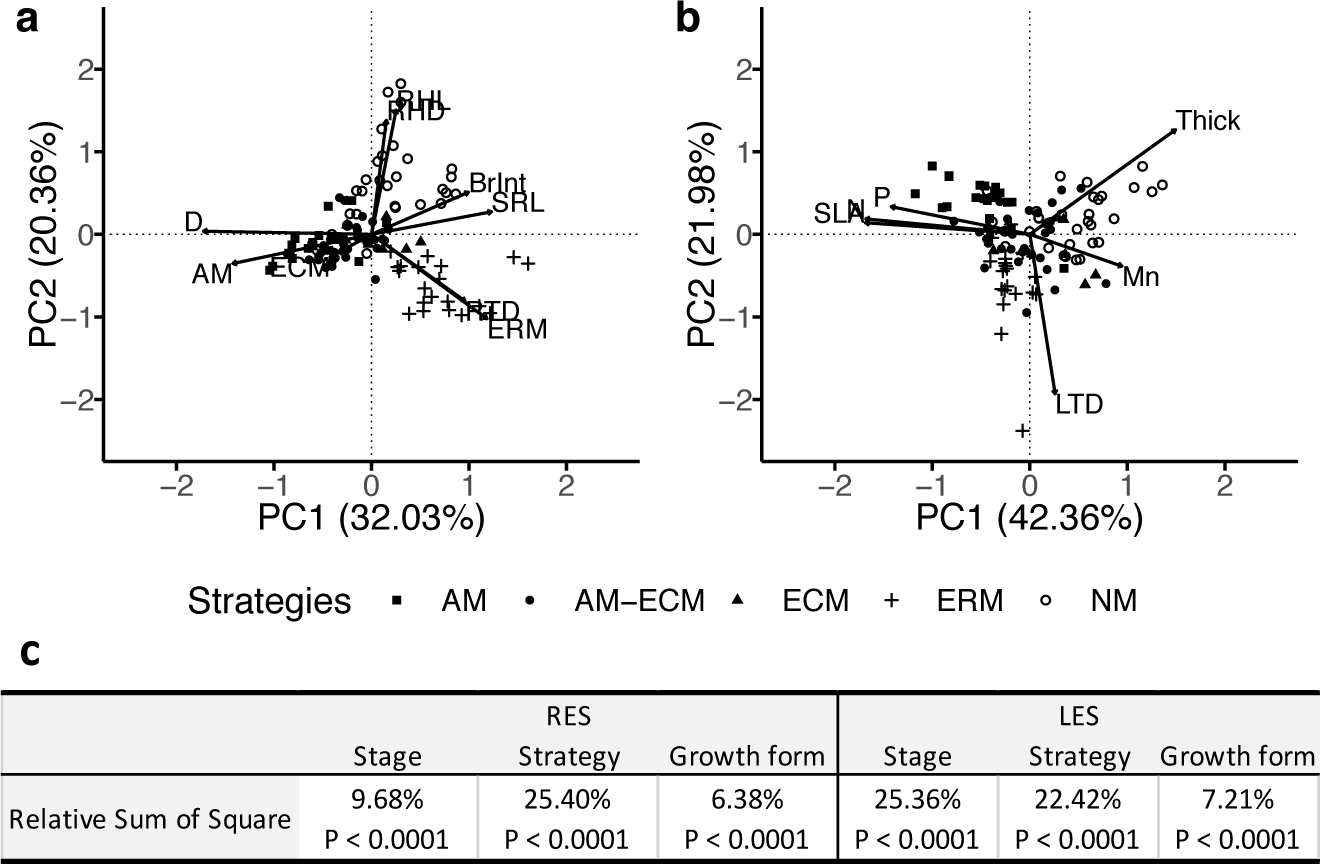
Projection of species in a Mahalanobis space formed by the two first principal components. (a) Principal component analysis performed on root functional traits; (b) performed on leaf functional traits; (c) distance-based analysis results. Abbreviations: SLA: specific leaf area, Thick: leaf thickness, P: leaf phosphorus concentration, N: leaf nitrogen concentration, Mn: leaf manganese concentration, LTD: leaf tissue density, RTD: root tissue density, SRL: specific root length, BrInt: branching intensity, D: root diameter, AM: arbuscular mycorrhizal colonization, ECM: ectomycorrhizal colonization, ERM: ericoid colonization, RHL: root hair length and RHD: root hair density. Strategy abbreviations: AM: arbuscular mycorrhizal host, ECM: ectomycorrhizal host, AM-ECM: dual arbuscular mycorrhizal and ectomycorrhizal host, ERM: ericoid mycorrhizal host and NM: non-mycorrhizal.

Similar to the root economics spectrum, PCA on leaf traits showed one major dimension, representing 42.4% of the variance, ranging from species with ‘fast’ leaves displaying high values of SLA, leaf [N] and [P] to species with ‘slow’ leaves exhibiting thick and dense leaves with high Mn concentration (Fig. 1b). Leaf tissue density contributed to an independent second dimension explaining 22.0% of the variance (Figs. 1b and S1b; Table S8). As observed in the root traits spectrum, this pattern of correlation was mainly driven by ericoid species, which had dense leaves (Figs. 1b and S2a).

PERMANOVA analysis revealed that both root and leaf trait spectra were mainly structured by nutrient-acquisition strategies (RES: relative sum of squares = 25.4%; LES: relative sum of squares = 22.4%) and chronosequence stage (RES: relative sum of squares = 9.7%; LES: relative sum of squares = 25.4%), and relatively weakly by growth forms (RES: relative sum of squares = 6.4%; LES: relative sum of squares = 7.2%). In both the RES and LES, all factors were significant at P < 0.0001 (Fig. 1c).

Fourth-corner analysis, which considers species abundance, revealed significant and strong correlations among root and leaf traits and soil fertility variables (Fig. 2a and b). ’Outsourcing’ root traits such as AM symbiosis (i.e., high AM colonization within abundant species) were associated with exchangeable inorganic P (resin P) (P = 0.052), soil total [P] (P = 0.031) and total [N] (P = 0.0027). High root diameter was also associated with high soil total [N] (P = 0.0027) and to a lesser extent with high resin P (P = 0.064). Root traits associated with an ‘autonomous’ strategy (i.e., high SRL, BrInt, and RHD) were weakly associated (0.05 < P < 0.1) with high soil total [P], total [N] and to resin [P]. High root hair length was the only morphological trait displaying a significant association with high soil total [P] (P = 0.044) (Fig. 2b). The only trait correlated with the conservation gradient, the RTD, although not significantly associated with soil descriptors (P > 0.05), displayed a trend in which a high RTD was associated with higher resin [P]. Leaf traits exhibiting a ‘fast’ strategy were all associated with high soil nutrient concentrations, as SLA was positively correlated with soil total [P] (P = 0.0018), total [N] (P = 0.0006) and resin [P] (P = 0.0031); leaf [P] was associated with high soil total [P] (P = 0.004), total [N] (P = 0.036), and to a lesser extent resin [P] (P = 0.061). High leaf [N] was associated with high soil total [N] (P = 0.0046) and total [P] (P = 0.0018). Finally, leaf thickness, which is representative of a ‘slow’ strategy, was associated with low soil total [P] (P = 0.0006) and total [N] (P = 0.0006) and resin [P] (P = 0.013), and high leaf [Mn] was associated with soil total [P] (P = 0.0027) and not with other soil descriptors (P > 0.05), and LTD was not significantly associated with any soil descriptors (P > 0.05) (Fig. 2a).

**Figure 2:**
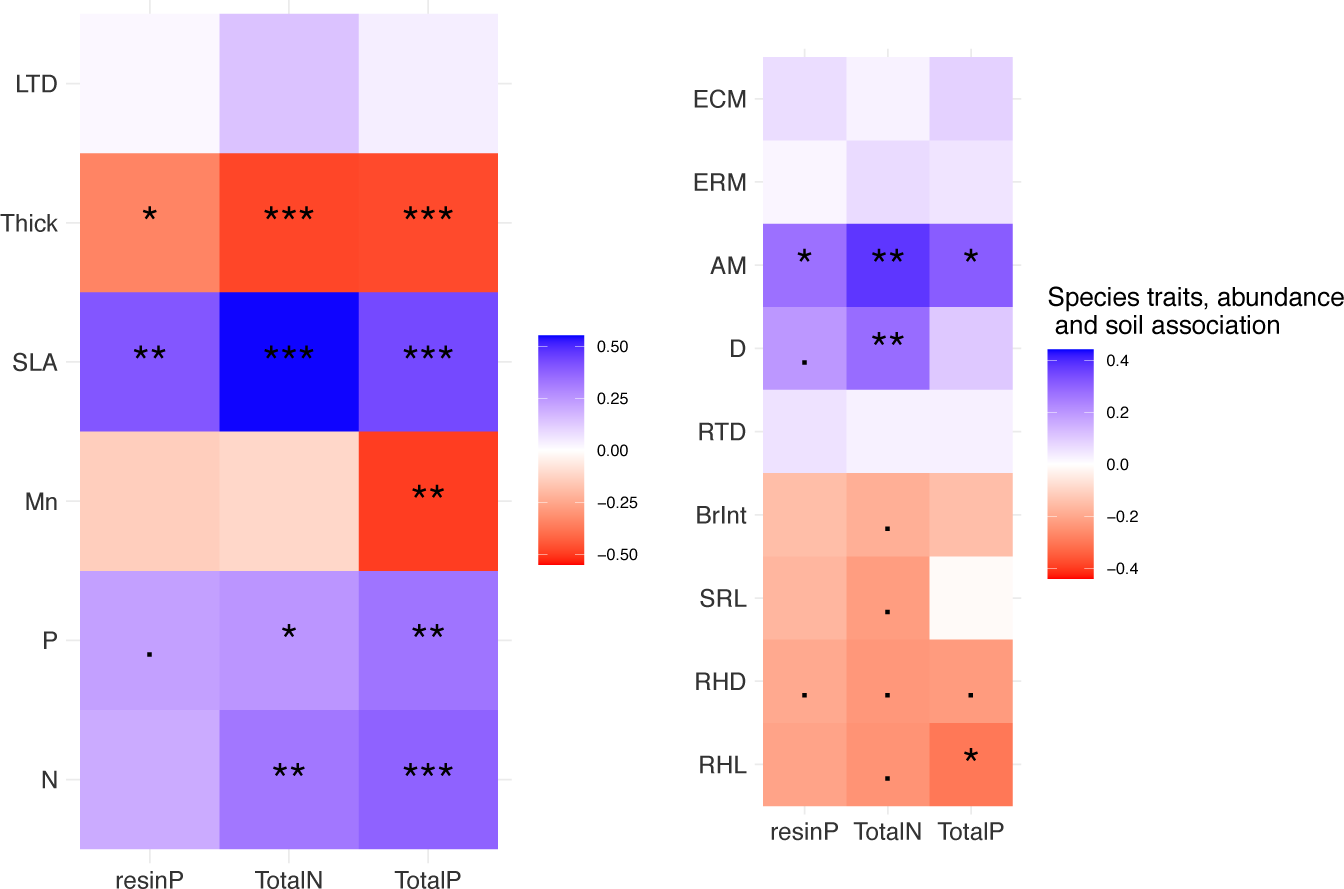
Fourth-corner analysis produced on the vegetation cover (Hellinger transformation), roots (a) and leaves (b) functional traits (normalized) within a cumulative relative cover of 75% and soil descriptors (normalized). The stars indicate the level of significance: (.) = P < 0.1, (*) = P < 0.05, (**) = P < 0.01 and (***) = P < 0.001. For abbreviations, see Figure 1

Community-weighted mean trait values and abundance of associated nutrient-acquisition strategies showed that stage-specific trait syndromes comprised multiple mycorrhizal strategies (Fig. 3a-e). However, irrespective of strategies, abundant species converged to thicker roots in soils displaying the highest nutrient concentration, with the exception of *Lepidosperma gladiatum* (Cyperaceae), a less common NM species. They also converged to longer root hairs or high SRL in nutrient-poor soils, with the exception of *Patersonia occidentalis*, a less common AM species and *Leucopogon obovatus* subsp*. revolutus*, a less common ERM species. Intraspecific variation in nutrient-acquisition strategy was substantial. Among the 15 species occurring in multiple plots, five of them exhibited different mycorrhizal status on at least two occasions (Table S3). This variation was further highlighted by the fact that each community comprised species displaying distinct root trait syndromes, as evidenced in this study by the stages with the most pronounced contrast in nutrient availability (Figs. S3; S4). Likewise, *Agonis flexuosa*, a common species occurring across all stages, differed in its mycorrhizal nutrient-acquisition strategies across stages (i.e., NM and ECM in stage 1, AM-ECM and ECM in stage 2, AM-ECM in stages 3 and 4 and AM-ECM and ECM in stage 5) (Table S3). Soil fertility-induced intraspecific variation was further evidenced by the fact that two of the most abundant species (i.e., *Agonis flexuosa* and *Bossiaea linophylla*) at the stage displaying the highest soil [N] and [P], although ECM at the youngest stage, converged to an AM-ECM strategy, as strictly ECM strategy was filtered out (Table S3; Fig. 3e). Finally, the impact of soil fertility on trait value variation was evidenced by the fact that the stage-specific leaf and root trait syndromes of common species *Agonis flexuosa* aligned with those of the entire plant community (Fig. 3b and d).

**Figure 3:**
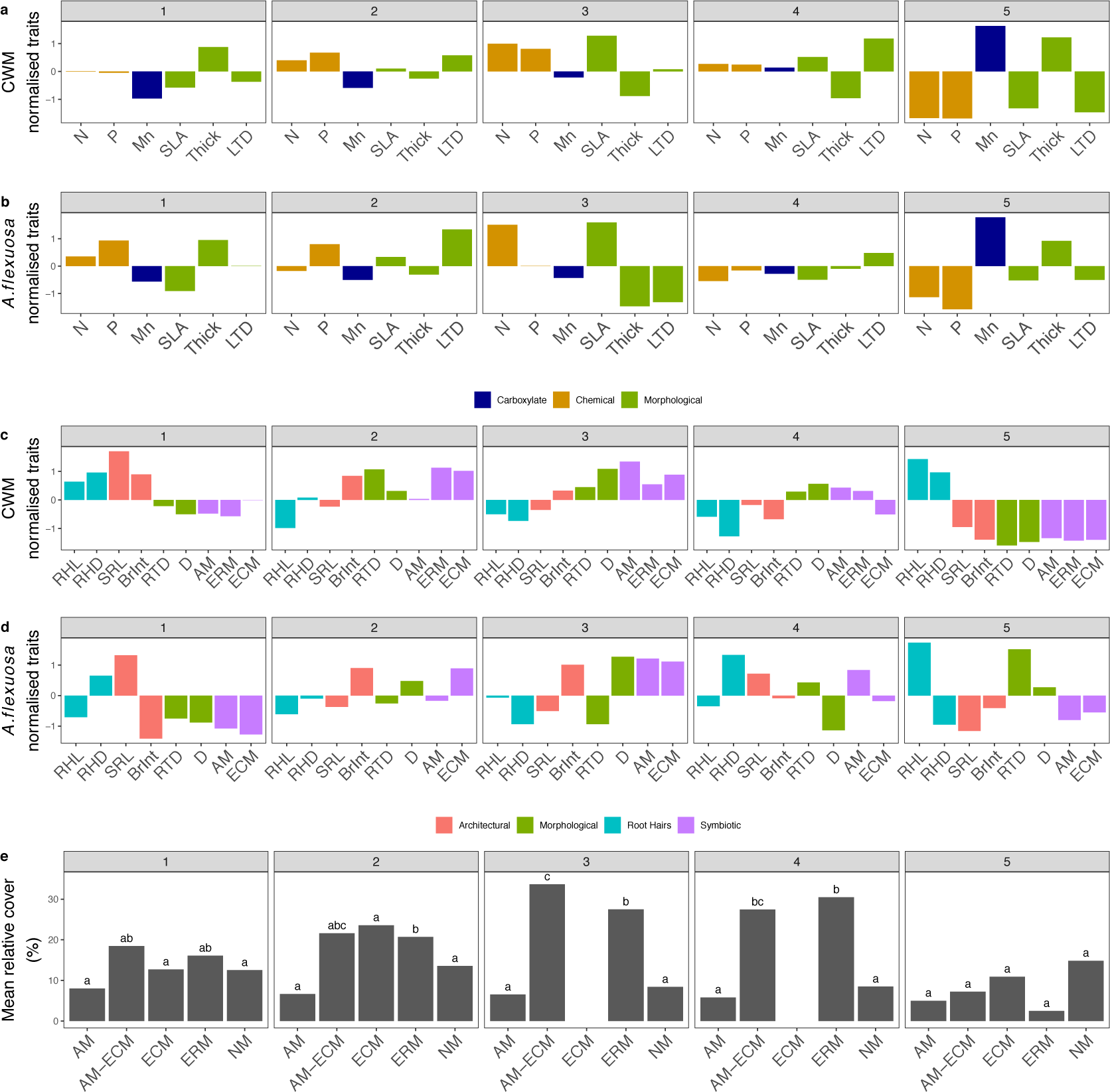
Community-weighted mean (CWM) common species traits and overall strategies mean relative cover (%) across chronosequence stages 1 to 5. a) CWM leaf traits; b) *Agonis flexuosa* leaf traits; c) CWM root traits; d) *Agonis flexuosa* root traits; and e) overall mean relative cover of mycorrhizal types across the chronosequence. No linear analyses were performed on CWM, since these analyses were performed in the fourth-corner analysis. Error bars in e) represent the 95% confidence intervals; letters above each mean represent Tukey honest significant difference (HSD) groupings (*P* ≤ 0.05). For strategy and for trait abbreviations, see Figure 1.

### Leaf and root traits coordination

Co-inertia analysis showed that leaf and root traits were weakly coordinated for abundance-unweighted traits (RV= 0.29; P < 0.001) and strongly coordinated for abundance-weighted traits (RV = 0.96; P < 0.008) (Fig. 4a and b). Similar to results of the fourth-corner analysis, co-inertia analysis performed on abundance-weighted traits revealed that ‘fast’ species traits within the LES covaried with ‘outsourcing’ species traits within the RES. Arbuscular mycorrhizal colonization, the pair ECM colonization and BrInt, the pair ERM colonization and RTD, root diameter, SLA, leaf [P] and [N] were superimposed in leaf and root traits covariance spectrums. ‘Slow’ species traits within the LES covaried with ‘autonomous’ species traits, as leaf thickness, RHD and RHL were also superimposed along the axis explaining more of the covariance (93.44%), while leaf [Mn] loaded onto both co-inertia axes (Fig. 4a and b).

**Figure 4:**
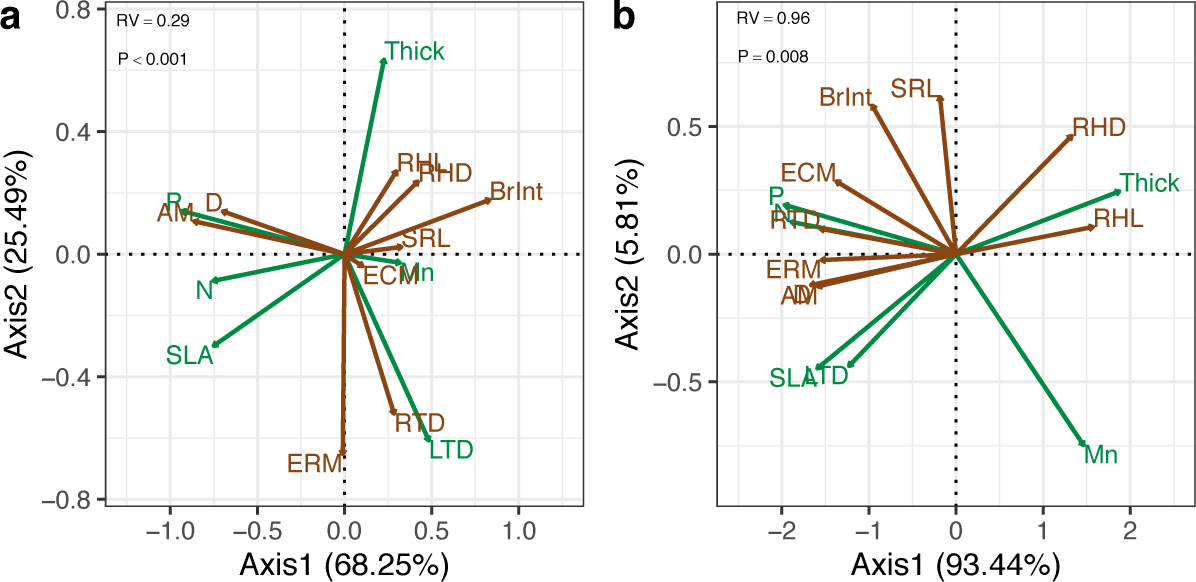
Leaf economic spectrum and root economic spectrum coinertia. *RV* coefficient is high when both structures vary along a similar axis, regardless of direction, and low when they vary independently. For unweighted root and leaf traits a) and for leaf and root traits weighted by relative cover. b). Green is associated with foliar traits and brown with root traits. For abbreviations, see Figure 1.

## Discussion

Mycorrhizal strategies structured the LES and RES, nevertheless, root and leaf traits were coordinated among each other across a strong soil P gradient once relative abundance was accounted for. Accounting for species abundance may have diluted the signal of rarer species that may acquire less abundant forms of nutrients, as observed in arctic plant communities for N (McKane et al., 2002) or tropical forests for P (Guilbeault-Mayers et al., 2020). Our results suggest that not only plant communities, but also common species whose leaves possess a ‘fast’ trait syndrome, meet their nutrient demands by coordinating with roots that exhibit a ‘collaborative’ trait syndrome in richer soils. On the other hand, in nutrient-impoverished soils, the ‘slow’ leaf trait syndrome was associated with an ‘autonomous’ root trait syndrome, which was characterized by longer root hairs and high leaf [Mn], pointing toward rhizospheric P mobilization through carboxylate exudation (Lambers et al., 2015a). Together, our results indicate that root traits defining the collaboration gradient responded to soil P availability and these traits were closely correlated with the aboveground ‘fast-slow’ leaf trait spectrum, which responded in the same manner. However, the species-level signal revealed that multiple nutrient-acquisition root trait syndromes might meet leaf demands along the LES suggesting that a single ‘fast-slow’ spectrum in an invalid concept to describe root and leaf trait coordination.

### Unconstrained interspecific variation in root and leaf traits

Our distance-based analysis results indicate that mycorrhizal nutrient-acquisition strategies, when used as a grouping factor, explained a larger proportion of the variance within the RES than soil fertility did, as also found by Valverde-Barrantes et al. (2017). Mycorrhizal association types may have played a significant role in explaining the observed variance, as a third significant dimension within the RES, contrasting ERM and ECM colonization, was observed. The contrasting investment in mycorrhizal association types might provide plants along this axis an increasing capacity to access recalcitrant and organic nutrients through the spectrum contrasting investment in ERM and ECM symbiosis (Ward et al., 2022). This suggests that the two dimensions of root trait variation identified by Bergmann et al. (2020) adequately capture inorganic nutrient-acquisition strategies but may not encompass strategies needed to access nutrient forms that are not readily available to fine roots or AM fungi. Meanwhile, nutrient-acquisition strategies explained slightly less variance than soil fertility within the LES which provides evidence that leaf trait values irrespective of mycorrhizal association type converged toward specific trait syndromes at all positions along the fertility gradient (Bernard-Verdier et al., 2012). We also found that growth forms explained a relatively small amount of variance within the RES and the LES, suggesting that a similar ecological trade-off structured variance, irrespective of growth form, for both LES (Wright et al., 2004) and RES (Bergmann et al., 2020). Overall, trait distribution within the LES and RES was primarily influenced by nutrient-acquisition strategies. Modification of nutrient-acquisition strategies in several common species along the chronosequence and the strong association between soil descriptors, species traits and abundance, however, support the idea that trait distribution reflected variation induced by soil fertility.

### Root and leaf trait variation constrained by a nutrient-availability gradient

Multivariate associations among soil fertility descriptors, species abundance and species traits revealed that all leaf traits, with the exception of LTD, were significantly associated with soil characteristics. Leaf traits linked with the ‘fast’ trait syndrome (i.e., high SLA, leaf [N] and [P]) were associated with more fertile soils and ‘slow’ trait values, represented by thick leaves were associated with nutrient-poor soils, as observed across and within biomes (Ordoñez et al., 2009; Jager et al., 2015). By contrast, among all root traits structuring the RES dimensions, only root traits associated with the ‘outsourcing’ trait syndrome (i.e., high AM colonization/root diameter) were significantly associated with high total [N], total [P] and exchangeable [P], while high values of root hair length combined with carboxylate-mobilizing P-mobilizing traits, proxied by high leaf [Mn] (Lambers et al., 2015a), were associated with extremely low soil total [P]. Although our results are not consistent with the trade-off underlying the collaboration gradient (i.e., diameter-SRL axis), they support a more comprehensive trade-off between root hair length and mycorrhizal symbioses that might effectively support rapid growth and survival at both ends of a nutrient-availability gradient. Long root hairs may enhance nutrient acquisition by expanding the root’s influence zone – including zones where neighboring plants’ carboxylate exudation might sporadically increase P availability – while matching the high leaf nutrient efficiency that supports survival in P-impoverished soil, given their resource-efficient nature (Lynch & Ho, 2005). Meanwhile, AM hyphae entail a lower carbon construction cost (Fitter, 1991), a higher P-acquisition rate per unit of length compared with fine roots (Jakobsen et al., 1992) and allow nutrient acquisition beyond the root’s nutrient-impoverished zone. AM colonization might, therefore, be conducive to growth in less infertile soil, provided that the potential increase in nutrient translocation to leaves (Eissenstat et al., 1993) offsets the higher respiration cost of colonized fine roots (Jakobsen & Rosendahl, 1990). The latter is supported by our study, as more abundant species displaying high diameter and AM colonization also exhibited high SLA, leaf [N], and [P], suggesting high photosynthetic activities (Wright et al., 2004). Overall, our results show that a tendency of high values of SRL, branching intensity, and root hair density – indicating an increase in nutrient foraging per unit length or volume – may be beneficial when nutrient availability is low. However, our study suggests a coordination between the LES and the collaboration gradient to support rapid growth in the most fertile soil and survival in nutrient-impoverished soil.

### Root and leaf traits coordination along a nutrient availability gradient

Co-inertia result confirm that the LES is strongly coordinated with the collaboration gradient – such as ‘outsourcing’ and ‘autonomous’ trait syndromes are positively associated with ‘fast’ and ‘slow’ leaf trait syndromes, respectively – when species abundance is accounted for. This aligns with the single ‘fast-slow’ spectrum, as an association between a trait syndrome along the diameter-SRL axis and specific soil nutrient content does not invalidate that this axis might be unrelated to variation in soil nutrient availability. Instead, it suggests that any trait syndrome along the diameter-SRL axis cannot be consistently associated with a given soil nutrient availability as proposed by (Guilbeault-Mayers & Laliberté, 2023). However, since the community-level RTD values showed a tendency for species to be positioned within the ‘slow’ RES subspace in the most fertile soil and in the ‘fast’ subspace in nutrient-impoverished soil, our results are not in line with a single ‘fast-slow’ spectrum (Ordoñez et al., 2009; Reich, 2014, Weigelt et al., 2021). Our results, however, align with Freschet et al.’s (2017), Liu et al.’s (2010), and Wen et al.’s (2022) proposition, which associates either rapid nutrient uptake for effective competition with microorganisms or rapid root exudation in nutrient-impoverished soil with the ‘fast’ end of the conservation gradient (i.e low RTD and high root [N]) which is generally required for these functions. The high RTD tendency observed in the most fertile soil (stage 3), however, was attributable to only two abundant species, *Banksia grandis* and *Leucopogon obovatus* subsp. *revolutus*. All other species within this stage, including two abundant AM-ECM species (*Agonis flexuosa* and *Bossiaea linophylla*), exhibited lower RTD values than the chronosequence mean (Fig. S3), thereby supporting a single ‘fast-slow’ spectrum. Likewise, our community-level results support that the ‘fast’ end of the conservation gradient was associated with P-impoverished soils (stage 5). Overall, species exhibited lower RTD values in stage 5, where soils are severely P-impoverished. However, a significant variation in RTD was observed at the species level. Notably, the common species *A. flexuosa* exhibited high RTD values in P-impoverished soil (Fig. S4). These latter two observations once again upheld the validity of both the single ‘fast-slow’ spectrum and a decoupling between ‘fast’ end of the LES, the ‘fast’ RES subspace and nutrient availability, depending on the scale of where the observations were made. Overall, this high local variation challenges the validity of a single ‘fast-slow’ spectrum at the whole plant scale.

### Challenging a single ‘fast-slow’ spectrum at the whole plant scale

The lack of convergence toward ‘fast’ or ‘slow’ root trait values among species calls into question the validity of a single ‘fast-slow’ spectrum (Reich, 2014). Stated differently, questioning its validity implies that a ‘fast-slow’ spectrum is shared between leaf and root functioning, but the ‘fast’ or ‘slow’ ends of these spectrums do not consistently align. That is, permanently categorizing fine roots as ‘slow’ or ‘fast’ solely based on individual trait values (Dallstream et al., 2022), or even trait syndromes may not be valid, contrasting with leaf functioning, for which such categorization is supported (Reich et al., 1997). However, a ‘fast’ leaf in terms of nutrient use requires effective acquisition and translocation of belowground resources (Reich, 2014) and this could be met using a diversity of root strategies and at different costs (Raven et al., 2018). That is, a ‘slower’ root strategy in a nutrient-richer soil, entailing, for example, faster metabolism and higher maintenance respiration costs, could be labeled as ‘fast’. This would hold true if that root strategy enabled the acquisition and translocation of adequate nutrients to meet the demands of a ‘fast’ leaf, and if the net carbon cost associated with this strategy allowed for sufficient carbon allocation to growth. For instance, within the stage where soil [N] and [P] were higher (stage 3), we observed both at the species level (i.e., five out of seven species) and at the community level ‘fast’ leaf trait syndromes. Nutrient acquisition to meet the demand of this leaf strategy, however, was achieved via high ERM colonization in *L. revolutus*, high AM colonization in *Hibbertia cuneiformis* and *H. grossulariifolia*, high AM-ECM colonization in *A. flexuosa* and long root hairs with low mycorrhizal colonization intensity in *B. linophylla* (Fig. S3). Furthermore, the latter two species have been recognized as carboxylate-dependent, as their nutrient uptake is partially reliant on P mobilization through carboxylate exudation by neighboring plants (Huang et al., 2017; Abrahão et al., 2018). Conversely, within the same stage, ‘slow’ leaf traits were displayed by *B. grandis* and *Lepidosperma gladiatum*, a Proteaceae and Cyperaceae, respectively. This might be accounted for by the higher overall cost associated with their carboxylate-releasing P-acquisition traits, which involve rapid root turnover (Lambers et al., 2015b) and rapid respiratory metabolism (Shane et al., 2004; Funayama-Noguchi et al., 2021).

## Conclusion

Overall, our results indicate that nutrient-acquisition strategies beyond the ‘fast’ RES subspace and even extending outside the RES plane as defined by Bergmann et al. (2020) into an axis opposing ERM and ECM colonization, as observed in the present study, might be labeled as ‘fast’. From this perspective, the resulting community’s abundance profile, which represents a species’ relative abundance, may arise from the selection of the most effective strategies during plant community assembly. This selection process, as suggested by Raven et al. (2018), might be influenced by competition for the same form of nutrient, resource partitioning (Turner, 2008), and facilitation of P acquisition, where one strategy may release a specific form of nutrient that becomes available to plants that exhibit another (Lambers et al., 2018). This suggests that leaf and root traits may exhibit local coordination within the ‘fast-slow’ spectrum framework. Considering that this spectrum is not unique, however, it is unlikely that a specific coordination pattern among leaf and root trait values might be consistently observed at a local scale across different biomes. However, given that root [N] was not measured, more studies including root exudation, leaf and root trait measurements along strong soil nutrient-availability gradients are needed to test if there is a single ‘fast-slow’ plant economic spectrum in which high root [N] might be consistently labeled as a ‘fast’ trait. Despite this, overall, the present study suggests that the same axes of fundamental trait variation are shared among leaf and root functioning, but that a single ‘fast-slow’ spectrum is an invalid concept to understand the correlation pattern among these axes across leaves and roots. Consequently, generalizing a specific set of root trait values that consistently meet the nutrient demands of a ‘fast’ or ‘slow’ leaf trait syndrome is unlikely at a local scale, and assuming this, global coordination among leaf and root traits might lead to uninformative results at smaller scales.

## Acknowledgements

Special thanks to Sharyn and Shaun Cody; without their kindness and advice, the project would not have been as successful. We also extend our gratitude to David Poissant and Caroline Fink-Mercier for their assistance in the field. Funding for this research was provided by a Discovery Grant from the Natural Sciences and Engineering Research Council of Canada (NSERC; grant RGPIN-2014-06106 and RGPIN-2019-04537). XGM received additional support from the Fonds de recherche du Québec-Nature et technologies.

## Author contributions

XGM, EL and HL conceived the study; XGM conceived the methodology; XGM conducted the data collection; XGM analyzed the data; XGM, HL and EL interpreted the results; XGM led the writing of the manuscript. All authors contributed to the draft versions.

## Data availability

The data that support the findings of this study will be openly available upon acceptance.

## References

Abrahão A, Ryan MH, Laliberté E, Oliveira RS, Lambers H. 2018. Phosphorus- and nitrogen-acquisition strategies in two *Bossiaea* species (Fabaceae) along retrogressive soil chronosequences in south-western Australia. Physiologia Plantarum 163(3): 323–343.

Bergmann J, Weigelt A, van der Plas F, Laughlin DC, Kuyper TW, Guerrero-Ramirez N, Valverde-Barrantes OJ, Bruelheide H, Freschet GT, Iversen CM et al. 2020. The fungal collaboration gradient dominates the root economics space in plants. Science Advances 6(27): eaba3756.

Bernard-Verdier M, Navas M-L, Vellend M, Violle C, Fayolle A, Garnier E. 2012. Community assembly along a soil depth gradient: Contrasting patterns of plant trait convergence and divergence in a Mediterranean rangeland. Journal of Ecology 100(6): 1422–1433.

Brundrett MC. 2009. Mycorrhizal associations and other means of nutrition of vascular plants: Understanding the global diversity of host plants by resolving conflicting information and developing reliable means of diagnosis. Plant and Soil 320(1–2): 37–77.

Claassen VP, Zasoski RJ. 1992. A containerized staining system for mycorrhizal roots. New Phytologist 121(1): 49–51.

Craine JM. 2009. Resource strategies of wild plants. Princeton, US: Princeton University Press.

Dallstream C, Weemstra M, Soper FM. 2022. A framework for fine-root trait syndromes: Syndrome coexistence may support phosphorus partitioning in tropical forests. Oikos 1: e08908.

Dray S, Chessel D, Thioulouse J. 2003. Co-inertia analysis and the linking of ecological data tables. Ecology 84(11): 3078–3089.

Dray S, Dufour A-B. 2007. The ade4 package: Implementing the duality diagram for ecologists. Journal of Statistical Software 22(4): 1–20.

Eissenstat DM, Graham JH, Syvertsen JP, Drouillard DL. 1993. Carbon economy of sour orange in relation to mycorrhizal colonization and phosphorus status. Annals of Botany, 71(1), 1–10.

Fitter AH. 1991. Costs and benefits of mycorrhizas: implications for functioning under natural conditions. Experientia 47: 350–355.

Freschet GT, Valverde-Barrantes OJ, Tucker CM, Craine JM, McCormack ML, Violle C, Fort F, Blackwood CB, Urban-Mead KR, Iversen CM et al. 2017. Climate, soil and plant functional types as drivers of global fine-root trait variation. Journal of Ecology 105(5): 1182–1196.

Funayama-Noguchi S, Shibata M, Noguchi K, Terashima I. 2021. Effects of root morphology, respiration and carboxylate exudation on carbon economy in two non-mycorrhizal lupines under phosphorus deficiency. Plant, Cell & Environment 44(2): 598–612.

Götzenberger L, de Bello F, Bråthen KA, Davison J, Dubuis A, Guisan A, Lepš J, Lindborg R, Moora M, Pärtel M et al. 2012. Ecological assembly rules in plant communities—approaches, patterns and prospects. Biological Reviews 87(1): 111–127.

Guilbeault-Mayers X, Laliberté E. 2023. Root phosphatase activity is coordinated with the root conservation gradient across a phosphorus gradient in a lowland tropical forest. bioRxiv.

Guilbeault-Mayers X, Turner BL, Laliberté E. 2020. Greater root phosphatase activity of tropical trees at low phosphorus despite strong variation among species. Ecology 101(8): e03090.

Hothorn T, Bretz F, Westfall P. 2008. Simultaneous inference in general parametric models. Biometrical Journal 50(3): 346–363.

Huang G, Hayes PE, Ryan MH, Pang J, Lambers H. 2017. Peppermint trees shift their phosphorus-acquisition strategy along a strong gradient of plant-available phosphorus by increasing their transpiration at very low phosphorus availability. Oecologia 185(3): 387–400.

Jager MM, Richardson SJ, Bellingham PJ, Clearwater MJ, Laughlin DC. 2015. Soil fertility induces coordinated responses of multiple independent functional traits. Journal of Ecology 103(2): 374–385.

Jakobsen I, Abbott LK, Robson AD. 1992. External hyphae of vesicular-arbuscular mycorrhizal fungi associated with *Trifolium subterraneum* L. New Phytologist 120(3): 371–380.

Jakobsen I, Rosendahl L. 1990. Carbon flow into soil and external hyphae from roots of mycorrhizal cucumber plants. New Phytologist 115(1): 77–83.

Kendrick GW, Wyrwoll KH, Szabo BJ. 1991. Pliocene-Pleistocene coastal events and history along the western margin of Australia. Quaternary Science Reviews 10(5): 419–439.

Laliberté E. 2017. Below-ground frontiers in trait-based plant ecology. New Phytologist 213(4): 1597– 1603.

Lambers H, Albornoz F, Kotula L, Laliberté E, Ranathunge K, Teste FP, Zemunik G. 2018. How belowground interactions contribute to the coexistence of mycorrhizal and non-mycorrhizal species in severely phosphorus-impoverished hyperdiverse ecosystems. Plant and Soil 424(1): 11– 33.

Lambers H, Hayes PE, Laliberté E, Oliveira RS, Turner BL. 2015a. Leaf manganese accumulation and phosphorus-acquisition efficiency. Trends in Plant Science 20(2): 83–90.

Lambers H, Martinoia E, Renton M. 2015b. Plant adaptations to severely phosphorus-impoverished soils. Current Opinion in Plant Biology 25: 23–31.

Lenth R. 2020. Emmeans: Estimated Marginal Means, aka Least-Squares Means. R package version 1.5.1. URL https://CRAN.R-project.org/package=emmeans.

Liu G, Freschet GT, Pan X, Cornelissen JHC, Li Y, Dong M. 2010. Coordinated variation in leaf and root traits across multiple spatial scales in Chinese semi-arid and arid ecosystems. New Phytologist 188(2): 543–553.

Lynch JP, Ho MD. 2005. Rhizoeconomics: Carbon costs of phosphorus acquisition. Plant and Soil 269(1–2): 45–56.

Ma Z, Guo D, Xu X, Lu M, Bardgett RD, Eissenstat DM, McCormack ML, Hedin LO. 2018. Evolutionary history resolves global organization of root functional traits. Nature 555(7694): 94– 97.

McKane RB, Johnson LC, Shaver GR, Nadelhoffer KJ, Rastetter EB, Fry B, Giblin AE, Kielland K, Kwiatkowski BL, Laundre JA et al. 2002. Resource-based niches provide a basis for plant species diversity and dominance in arctic tundra. Nature 415(6867): 68–71.

Oksanen J, Blanchet FG, Friendly M, Kindt R, Legendre P, McGlinn D, Minchin PR, O’Hara RB, Simpson GL, Solymos P et al. 2020. vegan: Community Ecology Package (Version 2.5-7). URL https://CRAN.R-project.org/package=vegan

Ordoñez JC, Van Bodegom PM, Witte J-PM, Wright IJ, Reich PB, Aerts R. 2009. A global study of relationships between leaf traits, climate and soil measures of nutrient fertility. Global Ecology and Biogeography 18(2): 137–149.

Peres-Neto PR, Jackson DA, Somers KM. 2003. Giving meaningful interpretation to ordination axes: Assessing loading significance in principal component analysis. Ecology 84(9): 2347–2363.

Pinheiro J, Bates D, Sarkar D, R Core Team. 2020. Nlme: Linear and nonlinear mixed effects models_. R package version 3.1–149. URL https://CRAN.R-project.org/package=nlme

Playford PE, Cockbain AE, Lowe GH. 1976. Geology of the Perth Basin, Western Australia; Bulletin of the Geological Survey of Western Australia. Geological Survey of Western Australia, Perth, Australia.

R Core Team. 2020. A language and environment for statistical computing. R Foundation for Statistical Computing, Vienna, Austria. URL https://www.R-project.org/.

Raven JA, Lambers H, Smith SE, Westoby M. 2018. Costs of acquiring phosphorus by vascular land plants: Patterns and implications for plant coexistence. New Phytologist 217(4): 1420–1427.

Reich PB. 2014. The world-wide ‘fast–slow’ plant economics spectrum: A traits manifesto. Journal of Ecology 102(2): 275–301.

Reich PB, Walters MB, Ellsworth DS. 1992. Leaf life-span in relation to leaf, plant, and stand characteristics among diverse ecosystems. Ecological Monographs 62(3): 365–392.

Reich PB, Walters MB, Ellsworth DS. 1997. From tropics to tundra: Global convergence in plant functioning. Proceedings of the National Academy of Sciences of the United States of America 94(25): 13730–13734.

Shane MW, Cramer MD, Funayama-Noguchi S, Cawthray GR, Millar AH, Day DA, Lambers H. 2004. Developmental physiology of cluster-root carboxylate synthesis and exudation in Harsh Hakea. Expression of phosphoenolpyruvate carboxylase and the alternative oxidase. Plant Physiology 135(1): 549–560.

Smith JE, Read DJ. 2008. Mycorrhizal symbiosis (Third Edition). New York, US: Academic Press.

Trouvelot A. 1986. Mesure du taux de mycorhization VA d’un systeme radiculaire. Recherche de methodes d’estimation ayant une significantion fonctionnelle. Mycorrhizae : Physiology and Genetics: 217–221.

Turner BL. 2008. Resource partitioning for soil phosphorus: A hypothesis. Journal of Ecology 96(4): 698–702.

Turner BL, Hayes PE, Laliberté E. 2018. A climosequence of chronosequences in southwestern Australia. European Journal of Soil Science 69(1): 69–85.

Turner BL, Romero TE. 2009. Short-term changes in extractable inorganic nutrients during storage of tropical rain forest soils. Soil Science Society of America Journal 73(6): 1972–1979.

Valverde-Barrantes OJ, Freschet GT, Roumet C, Blackwood CB. 2017. A worldview of root traits: The influence of ancestry, growth form, climate and mycorrhizal association on the functional trait variation of fine-root tissues in seed plants. New Phytologist 215(4): 1562–1573.

Vierheilig H, Coughlan AP, Wyss U, Piché Y. 1998. Ink and vinegar, a simple staining technique for arbuscular-mycorrhizal fungi. Applied and Environmental Microbiology 64(12): 5004–5007.

Walker LR, Wardle DA, Bardgett RD, Clarkson BD. 2010. The use of chronosequences in studies of ecological succession and soil development. Journal of Ecology 98(4): 725–736.

Walker TW, Syers JK. 1976. The fate of phosphorus during pedogenesis. Geoderma 15(1): 1–19.

Ward EB, Duguid MC, Kuebbing SE, Lendemer JC, Bradford MA. 2022. The functional role of ericoid mycorrhizal plants and fungi on carbon and nitrogen dynamics in forests. New Phytologist 235(5): 1701–1718.

Weemstra M, Mommer L, Visser EJW, van Ruijven J, Kuyper TW, Mohren GMJ, Sterck FJ. 2016. Towards a multidimensional root trait framework: A tree root review. New Phytologist 211(4): 1159–1169.

Weigelt A, Mommer L, Andraczek K, Iversen CM, Bergmann J, Bruelheide H, Fan Y, Freschet GT, Guerrero-Ramírez NR, Kattge J, et al. 2021. An integrated framework of plant form and function: The belowground perspective. New Phytologist 232(1): 42–59.

Wen Z, White PJ, Shen J, Lambers H. 2022. Linking root exudation to belowground economic traits for resource acquisition. New Phytologist 233(4): 1620–1635.

Wright IJ, Reich PB, Westoby M, Ackerly DD, Baruch Z, Bongers F, Cavender-Bares J, Chapin T, Cornelissen JH, Diemer M et al. 2004. The worldwide leaf economics spectrum. Nature 428(6985): 821–827.

Zuur AF, Ieno EN, Elphick CS. 2010. A protocol for data exploration to avoid common statistical problems. Methods in Ecology and Evolution 1(1): 3–14.

Zuur AF, Ieno EN, Walker N, Saveliev AA, Smith GM. 2009. Mixed effects models and extensions in ecology with R. New York, NY: Springer

