## Supporting information for "Coordination among leaf and fine root traits across a strong natural soil fertility gradient"

Table S1: Associations between root traits defining the collaboration gradient (i.e. diameter and specific root length) and soil fertility among studies. Positive signs (+) indicate that high values of this trait were associated with high nutrient availability, while negative signs (-) indicate that low values of this trait were associated with the same condition.

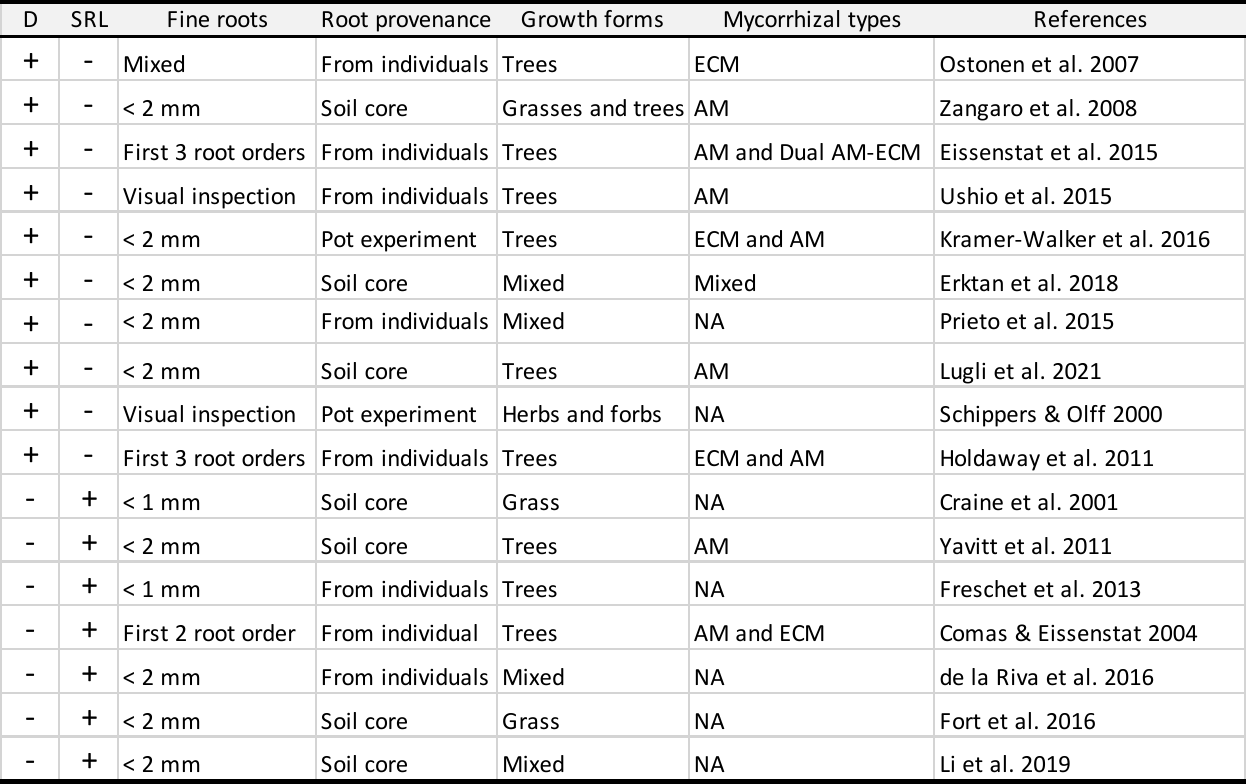

Table S2: Mean and range of soil characteristics across the chronosequence.

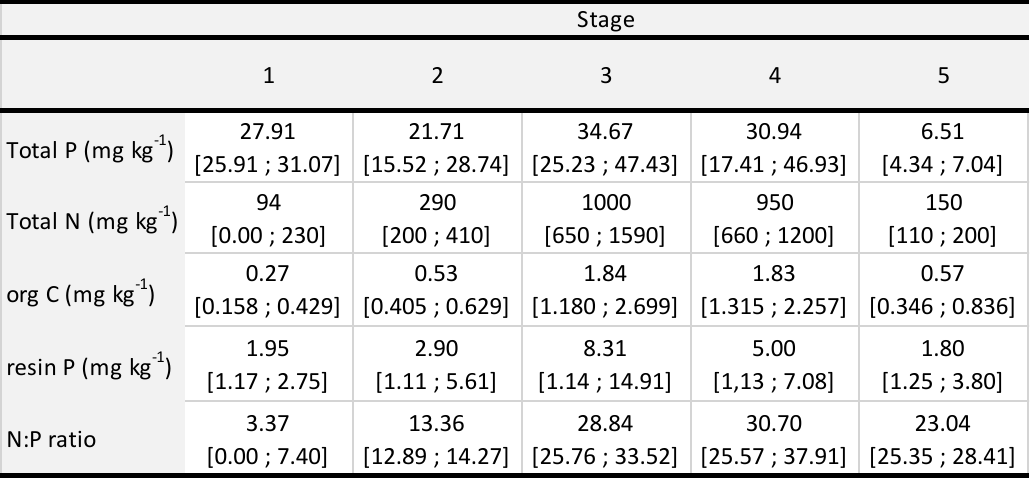

Table S3: Species sampled across the Warren chronosequence, showing for each species the mean and range of relative cover (%) per chronosequence stage. The last line of the table shows the mean and range of the cumulative relative cover of species within each chronosequence stage. The last two columns show the total number of composite samples taken and the change in mycorrhizal strategy. For strategy abbreviations, see figure S2.

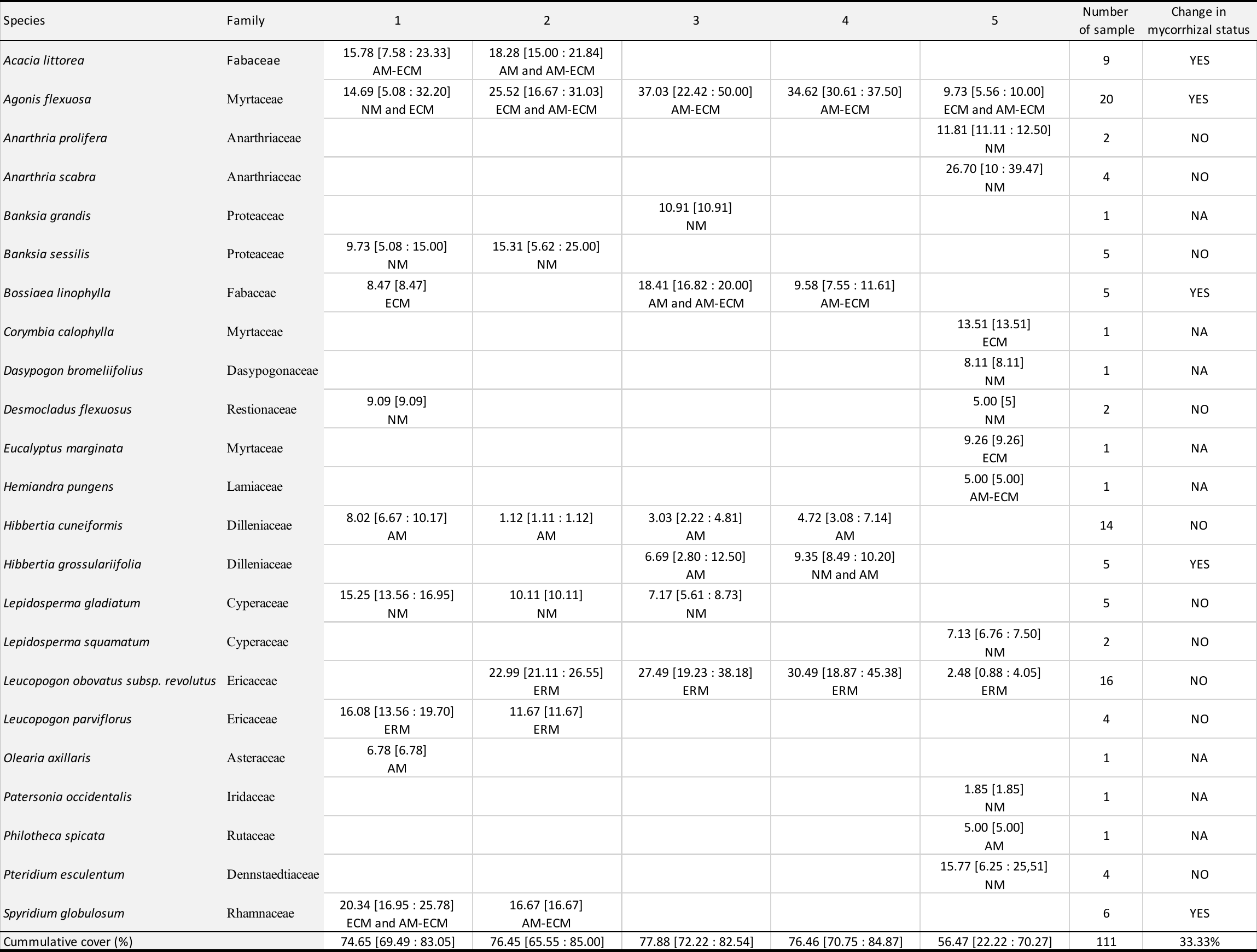

Table S4: Strategies of species sampled across the Warren chronosequence, showing for each strategy the absolute count and mean relative cover (%) per chronosequence stage. For abbreviations, see figure S2.

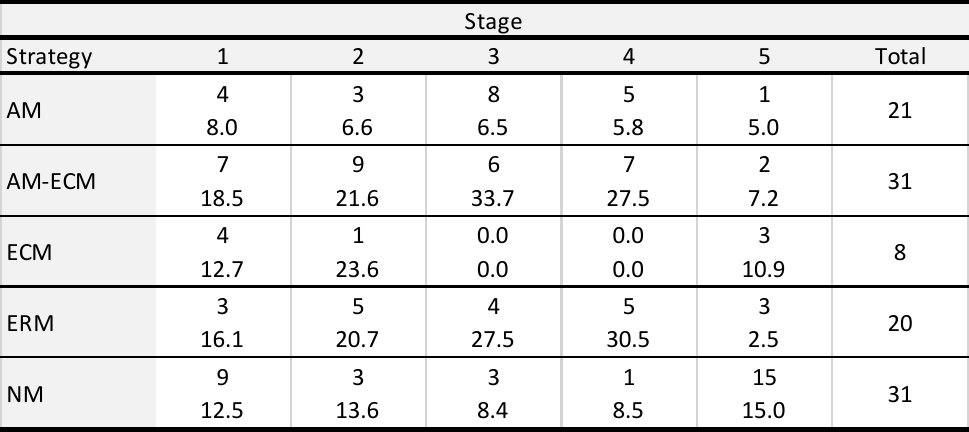

Table S5: Growth forms sampled across the Warren chronosequence, showing for each growth form the absolute count and mean relative cover (%) per chronosequence stage.

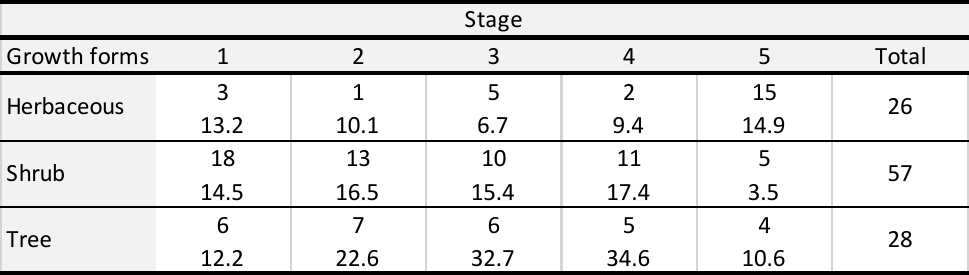

Table S6: Contingence table of species growth forms and strategies. For abbreviations, see figure S2.
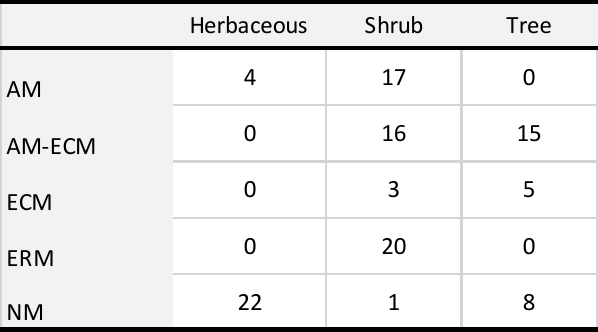

Table S7: Significant root trait loadings in principal component analysis. Trait loadings in **bold** explain more variance than the broken sticks distribution. For abbreviations, see figure S2.

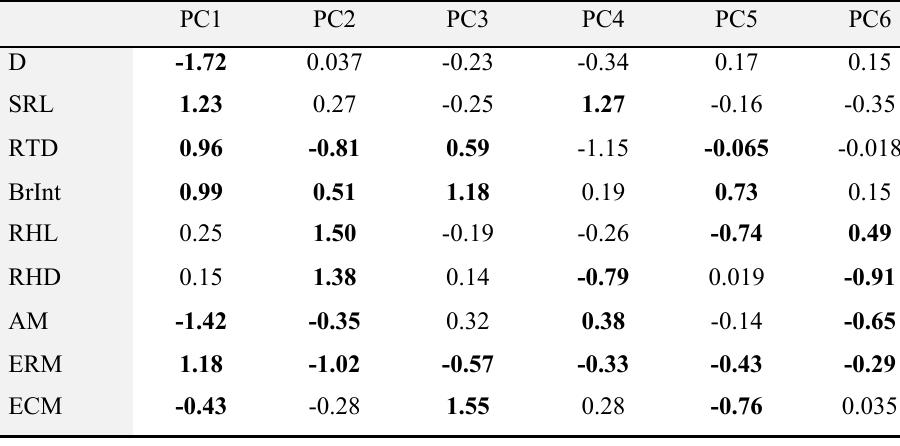

Table S8: Significant leaf trait loadings in principal component analysis. Trait loadings in **bold** explain more variance than the broken sticks distribution. For abbreviations, see Figure S2.

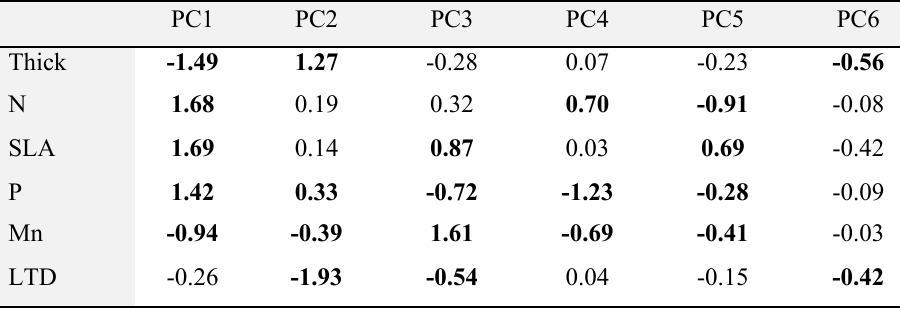

*Figure legends*

Figure S1: Dual representation of principal component analysis individual component inertia performed on observed and permuted data. Blue lines correspond to inertia axis values between the 5% and 95% quantiles based on 10,000 permutations and red to the observed inertia. Panel a) root traits and panel b) leaf traits.

Figure S2: Overall mean a) leaf trait and b) root trait values by strategy. Strategy abbreviations: AM: arbuscular mycorrhizal association, ECM: ectomycorrhizal association, ERM: ericoid association and NM: non-mycorrhizal. Error bars represent normalized standard deviation. Trait abbreviations: SLA: specific leaf area, Thick: leaf thickness, P: leaf phosphorus concentration, N: leaf nitrogen concentration, Mn: leaf manganese concentration, LTD: leaf tissue density, RTD: root tissue density, SRL: specific root length, BrInt: branching intensity, D: root diameter, AM: arbuscular mycorrhizal colonization, ECM: ectomycorrhizal colonization, ERM: ericoid colonization, RHL: root hair length and RHD: root hair density. Strategy abbreviations: AM: arbuscular mycorrhizal host, ECM: ectomycorrhizal host, AM-ECM: dual arbuscular mycorrhizal and ectomycorrhizal host, ERM: ericoid mycorrhizal host and NM: non-mycorrhizal.

Figure S3: Normalized root and leaf traits of aggregated stage 3 samples. In figures, 0 represents the overall chronosequence mean. For abbreviations, see Figure S2.

Figure S4: Normalized root and leaf traits of aggregated stage 5 samples. In figures, 0 represents the overall chronosequence mean. For abbreviations, see Figure S2.

Figure S1

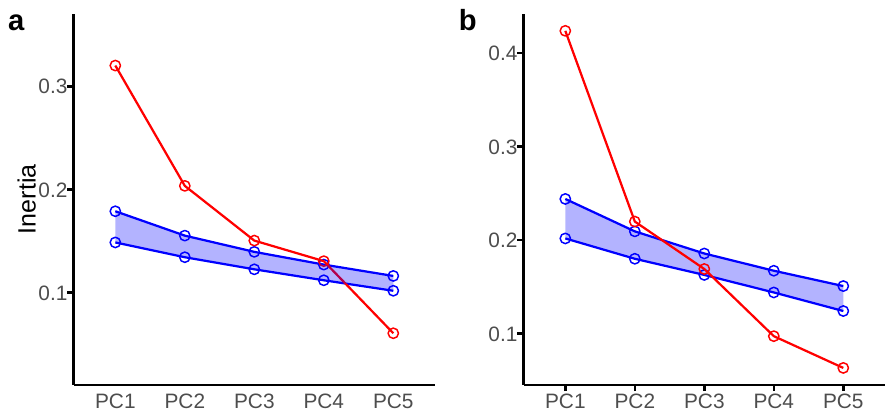

Figure S2

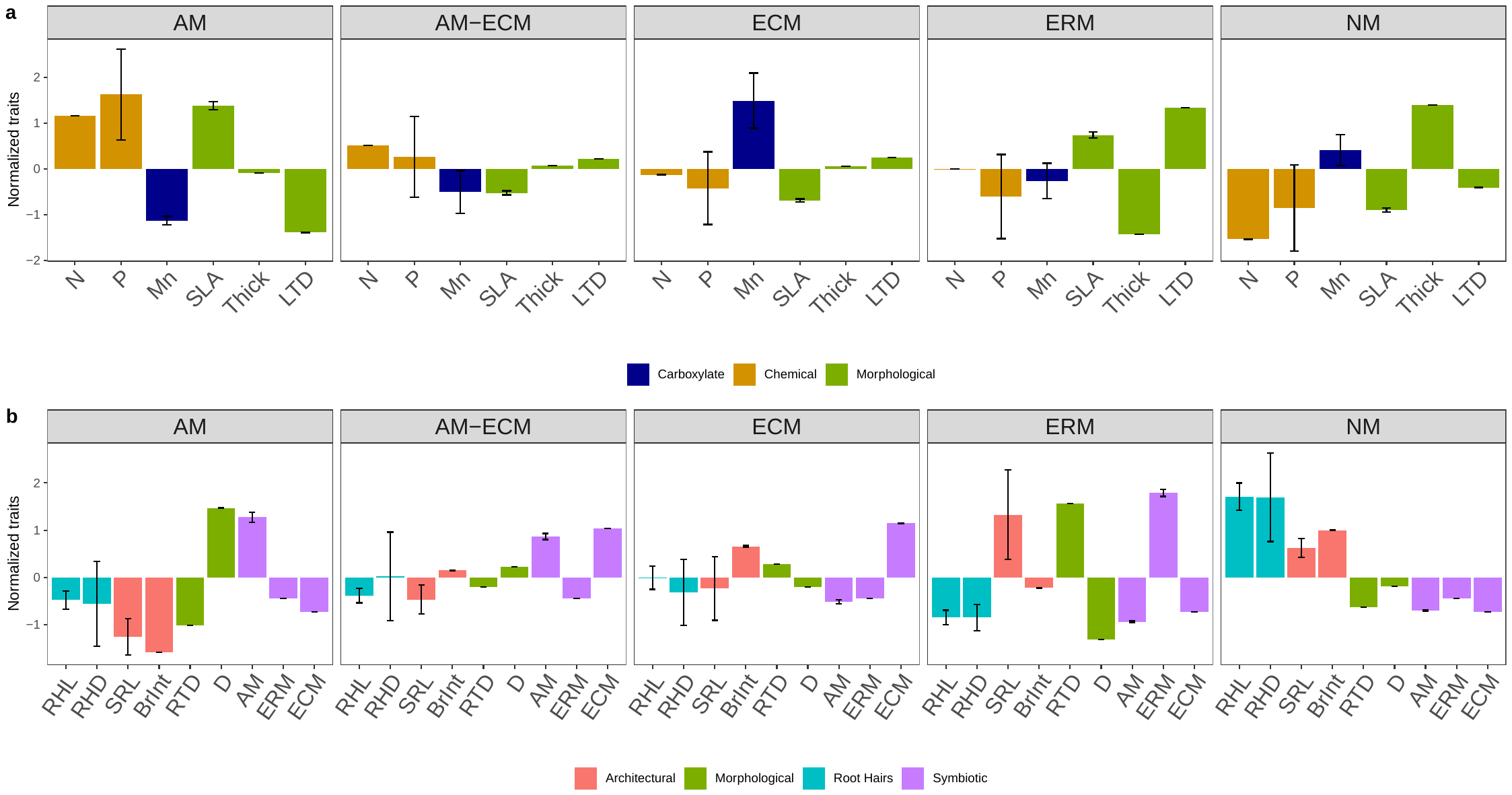

Figure S3

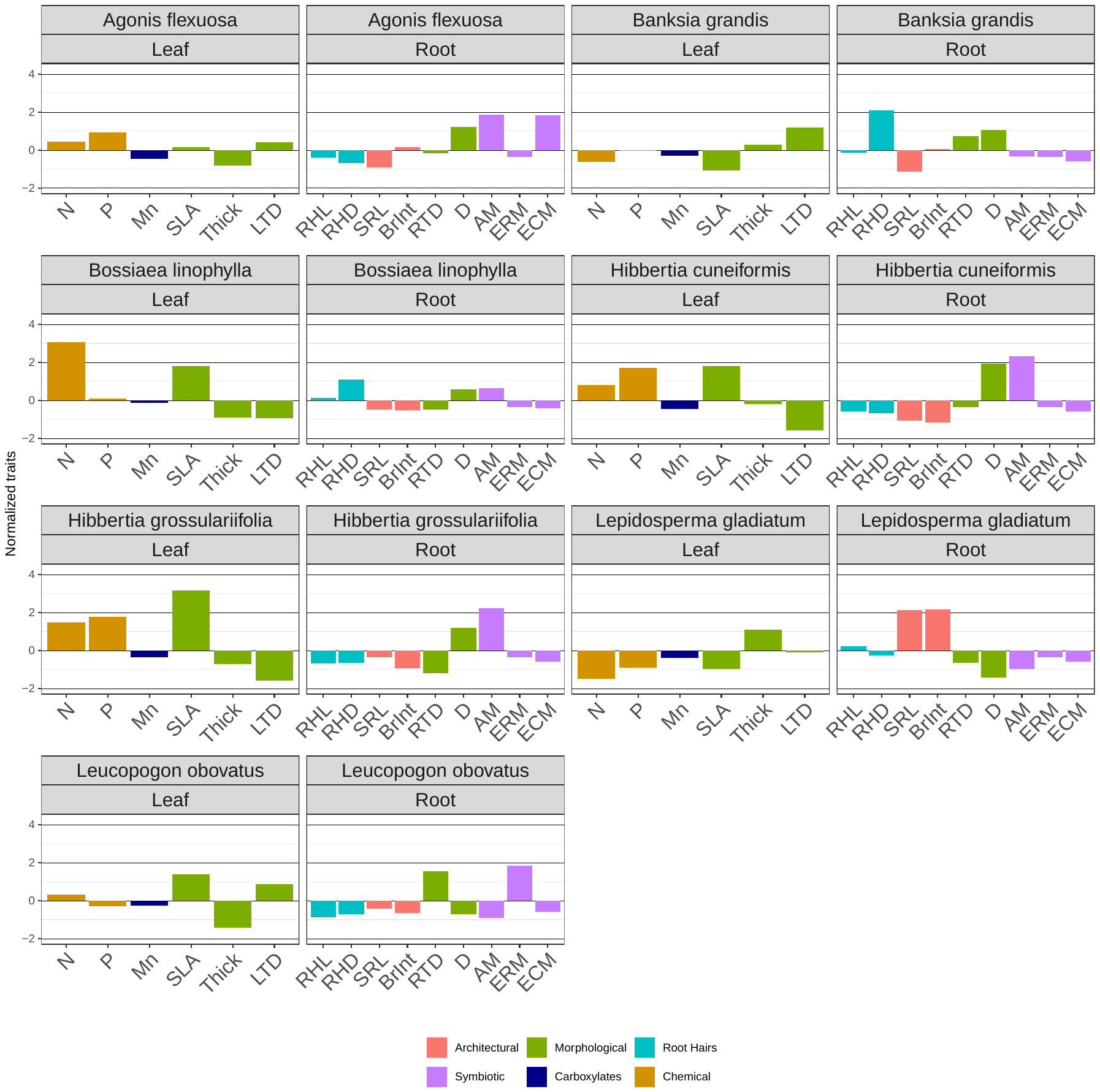

Figure S4

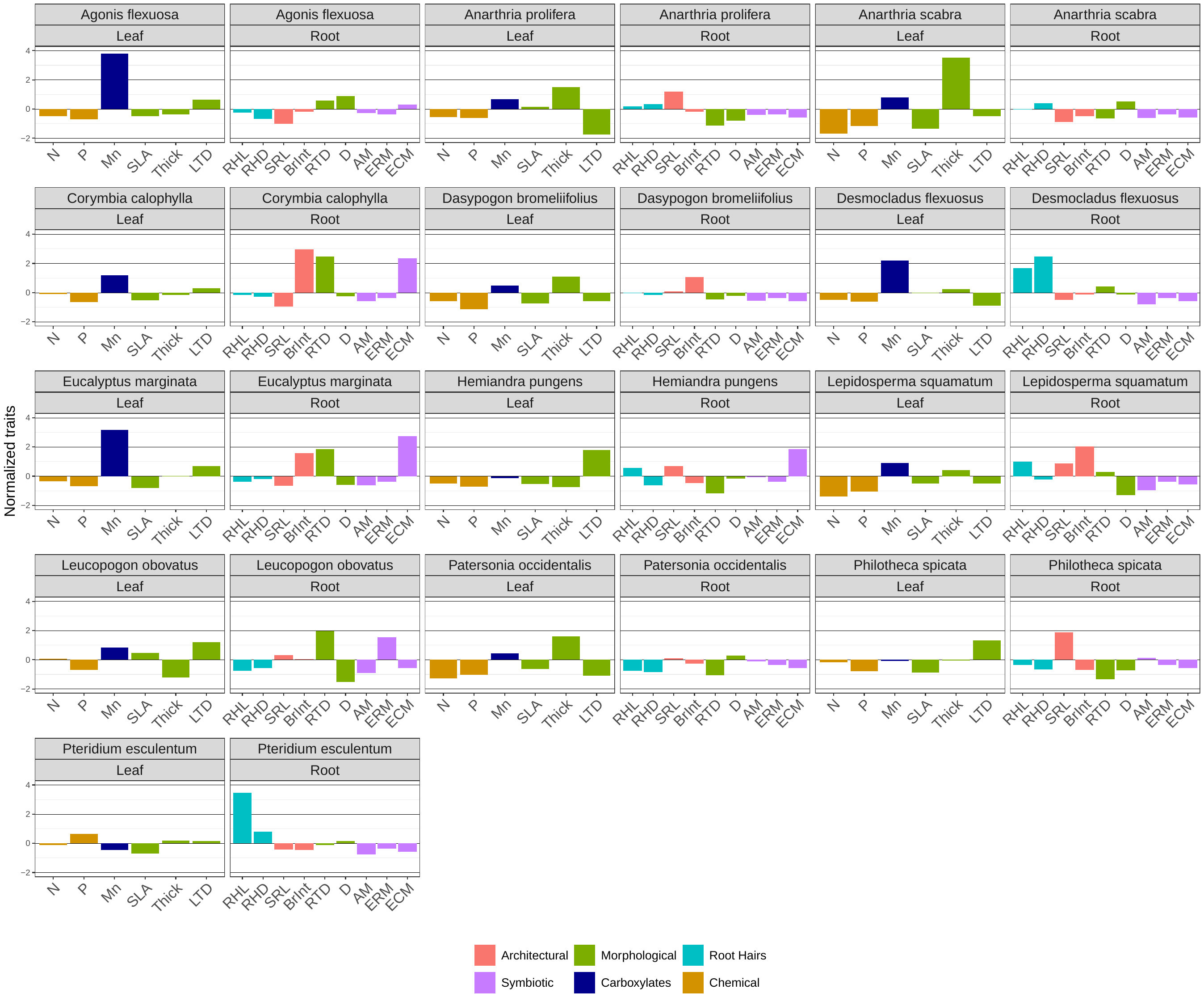

Table S1 references

Comas LH, Eissenstat DM. 2004. Linking fine root traits to maximum potential growth rate among 11 mature temperate tree species. Functional Ecology 18(3): 388–397.

Craine JM, Froehle J, Tilman DG, Wedin DA, Chapin FS. 2001. The relationships among root and leaf traits of 76 grassland species and relative abundance along fertility and disturbance gradients. Oikos 93(2): 274–285.

de la Riva EG, Tosto A, Pérez‐Ramos IM, Navarro‐Fernández CM, Olmo M, Anten NP, Marañón T, Villar R. 2016. A plant economics spectrum in Mediterranean forests along environmental gradients: Is there coordination among leaf, stem and root traits? Journal of Vegetation Science 27(1): 187–199.

Eissenstat DM, Kucharski JM, Zadworny M, Adams TS, Koide RT. 2015. Linking root traits to nutrient foraging in arbuscular mycorrhizal trees in a temperate forest. New Phytologist 208(1): 114–124.

Erktan A, Roumet C, Bouchet D, Stokes A, Pailler F, Munoz F. 2018. Two dimensions define the variation of fine root traits across plant communities under the joint influence of ecological succession and annual mowing. Journal of Ecology 106(5): 2031–2042.

Fort F, Cruz P, Lecloux E, Bittencourt de Oliveira L, Stroia C, Theau J-P, Jouany C. 2016. Grassland root functional parameters vary according to a community-level resource acquisition–conservation trade-off. Journal of Vegetation Science 27(4): 749–758.

Freschet GT, Bellingham PJ, Lyver PO, Bonner KI, Wardle DA. 2013. Plasticity in above- and belowground resource acquisition traits in response to single and multiple environmental factors in three tree species. Ecology and Evolution 3(4): 1065–1078.

Holdaway RJ, Richardson SJ, Dickie IA, Peltzer DA, Coomes DA. 2011. Species- and community-level patterns in fine root traits along a 120,000-year soil chronosequence in temperate rain forest. Journal of Ecology 99(4): 954–963.

Kramer-Walter KR, Bellingham PJ, Millar TR, Smissen RD, Richardson SJ, Laughlin DC. 2016. Root traits are multidimensional: Specific root length is independent from root tissue density and the plant economic spectrum. Journal of Ecology 104(5): 1299–1310.

Li F, Hu H, McCormack ML, Feng DF, Liu X, Bao W. 2019. Community-level economics spectrum of fine-roots driven by nutrient limitations in subalpine forests. Journal of Ecology 107(3): 1238–1249.

Lugli LF, Rosa JS, Andersen KM, Di Ponzio R, Almeida RV, Pires M, Cordeiro AL, Cunha HFV, Martins NP, Assis RL et al. 2021. Rapid responses of root traits and productivity to phosphorus and cation additions in a tropical lowland forest in Amazonia. New Phytologist 230(1): 116–128.

Ostonen I, Püttsepp Ü, Biel C. Alberton O, Bakker MR, Lõhmus K, Majdi H, Metcalfe D, Olsthoorn AFM, Pronk A et al. 2007. Specific root length as an indicator of environmental change. Plant Biosystems 141(3): 426–442.

Prieto I, Roumet C, Cardinael R, Dupraz C, Jourdan C, Kim JH, Maeght JL, Mao Z, Pierret A, Portillo et al. 2015. Root functional parameters along a land-use gradient: Evidence of a community-level economics spectrum. Journal of Ecology 103(2): 361–373.

Schippers P, Olff H. 2000. Biomass partitioning, architecture and turnover of six herbaceous species from habitats with different nutrient supply. Plant Ecology 149(2): 219–231.

Ushio M, Fujiki Y, Hidaka A, Kitayama K. 2015. Linkage of root physiology and morphology as an adaptation to soil phosphorus impoverishment in tropical montane forests. Functional Ecology 29(9): 1235-1245.

Yavitt JB, Harms KE, Garcia MN, Mirabello MJ, Wright SJ. 2011. Soil fertility and fine root dynamics in response to 4 years of nutrient (N, P, K) fertilization in a lowland tropical moist forest, Panama. Austral Ecology 36(4): 433–445.

Zangaro W, de Assis RL, Rostirola LV, de Souza PB, Gonçalves MC, Andrade G, Nogueira MA. 2008. Changes in arbuscular mycorrhizal associations and fine root traits in sites under different plant successional phases in southern Brazil. Mycorrhiza19(1): 37–45.
